# *E. coli* populations in unpredictably fluctuating environments evolve to face novel stresses through enhanced efflux activity

**DOI:** 10.1101/011007

**Authors:** Shraddha Madhav Karve, Sachit Daniel, Yashraj Deepak Chavhan, Abhishek Anand, Somendra Singh Kharola, Sutirth Dey

**Author notes:** Correspondence to, Sutirth Dey, Biology Division, Indian Institute of Science Education and Research Pune, Dr Homi Bhabha Road, Pune, Maharashtra, India 411 008. Present address: National Centre for Biological Sciences, GKVK, Bellary Road, Bangalore 560065, India.

## Abstract

There is considerable understanding about how laboratory populations respond to predictable (constant or deteriorating-environment) selection for single environmental variables like temperature or pH. However, such insights may not apply when selection environments comprise multiple variables that fluctuate unpredictably, as is common in nature. To address this issue, we grew replicate laboratory populations of *E. coli* in nutrient broth whose pH and concentrations of salt (NaCl) and hydrogen peroxide (H_2_O_2_) were randomly changed daily. After ∼170 generations, the fitness of the selected populations had not increased in any of the three selection environments. However, these selected populations had significantly greater fitness in four novel environments which have no known fitness-correlation with tolerance to pH, NaCl or H_2_O_2_. Interestingly, contrary to expectations, hypermutators did not evolve. Instead, the selected populations evolved an increased ability for energy dependent efflux activity that might enable them to throw out toxins, including antibiotics, from the cell at a faster rate. This provides an alternate mechanism for how evolvability can evolve in bacteria and potentially lead to broad spectrum antibiotic resistance, even in the absence of prior antibiotic exposure. Given that environmental variability is increasing in nature, this might have serious consequences for public-health.

## Introduction

In nature, organisms often face environments that fluctuate predictably or unpredictably. Such fluctuations are thought to affect a number of ecological and evolutionary processes including species coexistence (reviewed in Chesson, 2000) and diversity in a community (Abele, 1976; Hiltunen *et al*., 2008), evolution of invasive species (reviewed in Lee & Gelembiuk, 2008) and co-evolution (Harrison *et al*., 2013). Not surprisingly therefore, the evolutionary effects of environmental fluctuations have been thoroughly investigated, both theoretically (Levins, 1968) and empirically (reviewed in Kassen, 2002; Hedrick, 2006), leading to a number of insights. Stable environments tend to favor specialists (i.e. organisms with small niche width) whereas fluctuating environments often lead to the evolution of generalists (i.e. organisms with large niche width) (Kassen, 2002). Furthermore, evolutionary outcomes may depend on the periodicity of the environmental fluctuations. For instance, environmental shifts within the life-time of an organism can favor phenotypic plasticity (Ancel, 1999; Meyers *et al*., 2005), while environments that change over several hundreds of generations, particularly in asexual populations, can lead to evolution through selective sweeps of large-effect mutations (Cohan, 2005). However, when the environment fluctuates over intermediate time scales that are not long enough to promote selective sweeps and yet not short enough to select for plasticity, the evolutionary outcomes become more difficult to predict and might depend more critically on the underlying genetic architecture (Chevin *et al*., 2010).

Two other factors that can potentially determine the evolution of populations under environmental fluctuations are the predictability of the fluctuations (Hughes *et al*., 2007; Alto *et al*., 2013) and the complexity of the environments (Barrett *et al*., 2005; Cooper & Lenski 2010). Although there are some empirical studies that explicitly compare evolution under predictable and unpredictable fluctuations (Turner & Elena, 2000; Hughes *et al*., 2007), they typically concentrate on changes in a single environmental variable. However, in nature, environmental fluctuations typically happen simultaneously along multiple conditions which can potentially lead to a very different evolutionary response from a case where each of these stresses act individually. Thus, it is not clear whether the insights gained from evolutionary studies on unpredictable fluctuations in one environmental variable are relevant to cases where there are simultaneous fluctuations in multiple environmental variables. Furthermore, most environmental factors like temperature and pH are expected to vary on continuous scales, thus creating a much larger set of combinations experienced by the populations, compared to the scenario when a few discrete environmental combinations are investigated (Barrett *et al*., 2005; Cooper & Lenski 2010). Finally, there has been little elucidation of the mechanisms underlying adaptation to unpredictably fluctuating environments.

Here, we subjected replicate populations of *E. coli* to complex (combination of multiple values of pH, salt and H_2_O_2_), randomly changing environments. After ∼170 generations, the selected populations did not have any overall fitness advantage over the controls in the individual components of selection environments (i.e. pH, salt and H_2_O_2_). Interestingly, the selected populations had greater fitness when faced with novel stresses that did not have any known correlation with the selection environments. This fitness advantage continued to exist (albeit at lower levels) even after subjecting the selected and control populations to further ∼50 generations of directional selection in the novel environments. Contrary to expectations from theoretical studies, the mutation rates of the selected populations did not increase significantly, and there was no evolution of hypermutators. The ability to face novel stresses was partly explained by the evolution of greater efflux activity in the selected populations.

## Materials and Methods

Figure 1 shows a schematic of the various experiments performed in this study, along with the chief results for each.

**Fig 1:**
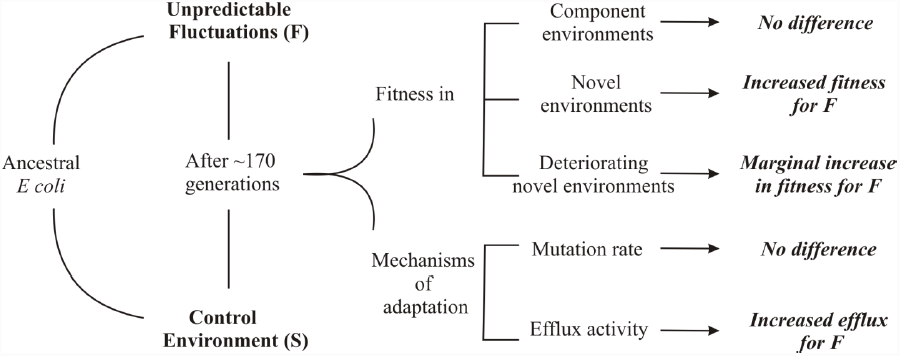
Schematic of the study along with the chief results for each experiment.

### 2.1 Selection experiments

#### 2.1.1 Selection under a stable environment

This study was conducted on *Escherichia coli* (strain NCIM 5547) populations. We plated this strain on nutrient agar, and created a suspension using bacteria from three randomly picked colonies. This suspension was used to initiate six bacterial cultures. Three of these were randomly assigned to a regime wherein they were grown at 37ºC, 150 rpm in 100 ml conical flasks. These three populations were sub-cultured every 24 hours (1ml in 50 ml of nutrient broth; see SOM-S1 for composition) for 30 days and will be henceforth referred to as the S (**S**table-Environment) populations. At the time of every sub-culturing, the optical density (OD_600_; measured using a Nanodrop^TM^, Thermo Scientific) was ∼0.2, signifying 5.64 generations every 24 hours (Bennett & Lenski, 1997). The other three populations were subjected to unpredictably changing complex environments for 30 days (see below) and will henceforth be referred as the F (**F**luctuating-Environment) populations. For both these sets of populations, we created glycerol stocks at the time of each sub-culturing.

#### 2.1.2 Selection under complex, randomly changing environments

In a pilot study, we observed the growth of this strain at various concentrations of salt (sodium chloride, NaCl), hydrogen peroxide (H_2_O_2_) and acidic / basic pH in nutrient broth (NB). These four conditions are henceforth referred to as *component environments*. Based on these observations, we created 72 arbitrary combinations of NaCl, pH and H_2_O_2_ values such that in any combination, the magnitude of one component was equal to that found in NB (i.e. pH = 7.0 or [NaCl] = 0.5g% or [H_2_O_2_] = 0) and that of the other two were individually expected to negatively affect growth. Thus, for example, combination # 46 denotes a NB containing 2 g% of NaCl + 9.5pH + no H_2_O_2_ whereas combination # 10 stands for 0.5 g% of NaCl + 9.5 pH + 0.01 M of H_2_O_2_ (SOM S1). Each combination was assigned a tag between 0-71 and a uniform-distribution random number generator was used to obtain a sequence of numbers (with replacement) in that range. Each replicate F population experienced 30 combinations of the components according to this sequence. This ensured that the environments faced by these populations were not only unpredictable, but also complex, i.e. involved three different environmental variables. As a result, unlike the S populations, the F populations did not grow to similar OD_600_ values in every generation. Therefore, we arbitrarily assigned an OD_600_ value ≥ 0.1 as a criterion for sub-culturing to the next environment. The volume used to inoculate the next culture was 1 ml, which translated to approximately 10^8^ −10^9^ cells. Any F population which did not reach an OD_600_ ≥ 0.1 was allowed to grow for another 24 hours in the same flask. If a population failed to cross the threshold even after 48 hrs post-inoculation, 100 µl of glycerol stock from the previous environmental combination was revived in NB and then transferred to the next combination in the selection sequence (pooling over all replicates, this event happened 20 times over the entire experiment).The total exposure to composite environments was kept equal, i.e. each replicate F population experienced 30 composite environments in which they could grow. All other culture conditions were identical to those experienced by the S populations.

### 2.2 Adaptation under different conditions

#### 2.2.1 Estimating Fitness

Following a recent study (Ketola et al., 2013b), we estimated fitness as the maximum growth rate during a defined period (here 24 hours). For all growth rate measurements mentioned below, the relevant glycerol stocks were revived in 50 ml nutrient broth for 18 to 20 hours. 4 µl of this revived culture (20 µl in case of selection under deteriorating environment) was inoculated into 2 ml of medium in 24-well cell culture plates and kept under 150 rpm at 37ºC. The OD_600_ of each well was measured every two hours over a period of 24 hours on a plate reader. The maximum growth rate of the bacterial population was then computed using a QBASIC (v 4.5) script to fit straight lines on overlapping moving windows of three points on the time series of OD_600_ values. The maximum slope obtained among all the fitted lines for a given time series, was taken as an estimate of the maximum growth rate for the corresponding population.

#### 2.2.2 Fitness under component environments

In this part of the study, we estimated the fitness of S and F populations (after ∼170 generations in their respective selection regimes) under four conditions of the component environments (i.e. salt, acidic pH, basic pH and H_2_O_2_).When a population is selected for resistance to a particular magnitude of stress (e.g. given concentration of antibiotic or a particular temperature) then adaptation can be inferred by assaying for fitness under the values of stress experienced during selection. However, in our study, the selection is characterized by unpredictable fluctuations in multiple stresses, over multiple values for each stress. Thus, it is not possible to designate a single value of the components as *the* selection environment. Therefore, we chose to assay the fitness of the F and S populations at the extreme values of the selected ranges for these environments (see Table S2 for concentrations used).

#### 2.2.3 Fitness under complex environments

In our study, what the selected populations actually experienced were combinations of stresses (i.e. pH+ salt +H_2_O_2_), which was designated as complex environments. Therefore, we also estimated the fitness of S and F populations (after ~170 generations in their respective selection regimes) under six randomly chosen complex environments faced by F populations during the selection (see Table S2 for details).

#### 2.2.4 Fitness under novel environments

In this part of the study, we assayed the fitness of the S and F populations (after ~170 generations in their respective selection regimes) under four different stressful conditions: presence of two heavy metals (cobalt and zinc), one antibiotic (norfloxacin) and a DNA intercalating agent (ethidium bromide). These four chemicals are known to reduce the growth rate of *E. coli*, and will be henceforth referred to as *novel environments*. The concentrations used in the assays are listed in Table S2. These four conditions were selected because we failed to find any study that reports a correlated change in fitness in any of them due to selection for any of the component environments (salt, pH or H_2_O_2_). Thus, it is unlikely that resistance to any of the four novel environments can evolve as a correlated response to direct selection on the component environments (see section 4.2 for discussion).

#### 2.2.5 Fitness after selection under deteriorating novel environments

We then investigated whether the fitness advantage of the selected populations over the controls in the novel environments, persisted even after directional selection. For this, all F and S populations were further selected separately under continuously increasing concentrations of cobalt, zinc and norfloxacin for 8 days (see Table S3 for details). Directional selection on ethidium bromide could not be done for logistical reasons. The maintenance protocol was the same as that for the selection under fluctuating environments (see section 2.1.2). After the 8^th^ day (i.e. ~ 50 generations), populations were stored as glycerol stocks for fitness estimation.

#### 2.2.6 Replicates and statistical analysis

For a given environment (e.g. cobalt or pH 4.5), the growth rate of each S and F population was measured in three replicates. All 18 growth rate measurements for a given environment (6 populations × 3 replicates) were conducted simultaneously in a single 24-well plate and constituted a trial. The entire process of revival and growth rate measurement for each component and novel environment was repeated one more time on a different date, to ascertain the consistency of the results. Thus, the study consists of two trials for fitness estimation in each environment, except under deteriorating novel environment, which consists of one trial.

To compare the fitness of the F and S populations averaged over multiple environmental conditions, we performed separate 4-way mixed-model ANOVAs for fitness under component environments, novel environments, and novel environments post directional selection. In each of these ANOVAs, selection (2 levels: S and F) and assay environments (3 or 4 levels) were treated as fixed factors crossed with each other. Replicate (3 levels, nested under selection) and trial (2 levels, nested in selection × assay environment × replicate), were treated as random factors. Since there was only one trial for the deteriorating environment, it was excluded from the analysis in that case. The fixed factor “assay environment” had four (acidic pH, basic pH, salt and H_2_O_2_), four (cobalt, zinc, norfloxacin and ethidium bromide) and three (cobalt, zinc, norfloxacin) levels in the ANOVAs on component environments, novel environments and novel environments post directional selection, respectively.

We were also interested in comparing the performance of the F and S populations in each assay-environment. Although, this can be done by comparing the appropriate means under the selection × assay environment interaction through planned-comparisons, we refrained from this approach. This is because such a comparison would neglect factors like replicates and trials, which were essential parts of our study design. Therefore, we followed a previous study (Ketola et al., 2013b) and performed separate 3-way mixed model ANOVAs for each assay environment and controlled for family-wise error rates through sequential Holm-Šidàk correction of the *p*-values (Abdi, 2010). This approach means that our interpretation, of a difference in fitness between the S and F populations being statistically significant, is conservative. The design of the ANOVAs was similar to the one above, except that assay environment was no longer a factor in the ANOVA.

In order to judge the biological significance of the differences in mean growth rates for F and S populations at different assay environments, we computed Cohen’s *d* statistic (Cohen, 1988) as a measure of the effect sizes (Sullivan & Feinn, 2012). Following existing guidelines (Cohen, 1988), we interpreted the effect sizes as small, medium and large for 0.2 < *d* < 0.5, 0.5 < *d* < 0.8 and *d* > 0.8 respectively.

All ANOVAs in this study were performed on STATISTICA v5.0 (Statsoft Inc.) while the Cohen’s *d*-statistics were estimated using the freeware Effect Size Generator v2.3.0 (Devilly, 2004).

### 2.3 Mechanism of adaptation

#### 2.3.1 Mutation rates

We used a well-established variant of fluctuation test (Crane et al., 1996) to estimate the mutation rates of F and S populations (see SOM – S4 for details). This method uses a median-based estimator (Jones, 1994) for the expected number of mutants and has been widely followed and recommended (Pope et al., 2008). The mutation-rate data were analyzed using a 1-way ANOVA with selection (2 levels: F and S) being a fixed factor.

#### 2.3.2 Efflux activity

One of the ways in which bacteria can fight multiple stresses is through active efflux of materials from the cell (Nies, 1999; Poole, 2005). A variant of an existing protocol (Webber & Coldham, 2010) was used to quantify the efflux activities of F and S populations (see SOM – S5 for details). The data were analyzed using a 2-way mixed model ANOVA where selection (2 levels: S and F) and replicates (3 levels, nested in selection) were treated as fixed and random factors respectively.

#### 2.3.3 Efflux activity under selection for adaptation to constant environment

In our study, any observed change in efflux activity of the F populations could be ascribed either to exposure to complex fluctuating environments, or to the effects of any of the individual component environments. To distinguish between these two possibilities, we subjected three replicate populations of *E. coli* each to selection for acidic pH, basic pH, salt and H_2_O_2_ (see SOM S6) for details of the selection process and the concentrations used). After ~40 generations, the efflux activities of these populations were measured and analyzed (see SOM S6 for the details).

## 3 Results

A non-technical summary of the major results is presented as figure 1.

### 3.1 Adaptation under different environmental conditions

#### 3.1.1 Fitness under component environments

After ∼170 generations of selection, the maximum growth rates of the F populations were significantly greater than the S, when measured under conditions that were components of the fluctuating selection regime, i.e. pH (4.5/10), high salt or H_2_O_2_ (*F_1,4_* = 9.34, *p* = 0.038, Table 1, Fig 2A). However, the effect size for this difference was small (*d* = 0.2, Table 1) which indicated that the difference might be biologically insignificant. There was also a significant main effect of component environment (F_3,12_ = 79.92, p = 3.4E-08) which is not surprising as the different conditions (i.e. pH 4.5, H_2_O_2_, etc.) are not expected to affect the growth rates similarly. However, the component environment × selection interaction was also significant (F_3,12_ = 3.850, *p* = 0.038), implying that there were differences in terms of how the growth rates of F and S populations were getting affected by the various conditions. To investigate this in greater detail, we conducted separate ANOVAs for the effect of each condition of the component environment and found that the growth rates of the F and S populations did not differ significantly in any of the stresses (Table 2, Fig 2A). The effect sizes of the differences were large for two conditions (pH4.5 and H_2_O_2_), medium for salt and small for pH10 (Table 2). Interestingly, in pH 4.5 although the difference has a large effect size, it is actually the S populations which have higher growth rates than the F. Overall, this set of analysis leads to the conclusion that in spite of a significant main effect, there is no evidence to indicate that the F populations generally performed better than the S under conditions that were components of the fluctuating selection regime.

**Table 1:** Summary of the main effects of selection in the pooled ANOVAs. Effect size was measured as Cohen’s *d* statistic and interpreted as small, medium and large for 0.2 < *d* < 0.5, 0.5 < *d* < 0.8 and *d* > 0.8 respectively.

| Assay | Mean S | Mean F | ANOVA F(1,4) | ANOVA p values | Effect Size±95% CI | Inference |
| --- | --- | --- | --- | --- | --- | --- |
| Fitness in Component Environments | 0.109 | 0.121 | 9.34 | 0.038 | 0.20±0.33 | Small |
| Fitness in Complex Environments | 0.074 | 0.088 | 2.37 | 0.199 | 0.34±0.27 | Small |
| Fitness in Novel Environments | 0.028 | 0.058 | 1320.48 | 3.4E-06 | 1.45±0.37 | Large |
| Fitness Post Deteriorating Environments | 0.076 | 0.086 | 24.68 | 0.008 | 0.51±0.54 | Medium |
| Mutation Rate | 1.0E-09 | 6.4E-09 | 4.11 | 0.11 | 1.65±1.85 | Large |
| Efflux potential | 0.204 | 0.364 | 34.65 | 0.004 | 3.79±1.55 | Large |

**Table 2:** Summary of the main effects of selection in the ANOVAs under individual environments. This table shows three set of fitness measurements, namely, component environments, novel environments and novel environments after directional selection. Effect size was measured as Cohen’s *d* statistic and interpreted as small, medium and large for 0.2 < *d* < 0.5, 0.5 < *d* < 0.8 and *d* > 0.8 respectively.

| Assay | Environment | Mean<br>S | Mean<br>F | ANOVA<br>F(1,4) | p values<br>(Holm-Šidák<br>corrected) | Effect<br>Size<br>±95% CI | Inference |
| --- | --- | --- | --- | --- | --- | --- | --- |
| Fitness in<br>component<br>environments | pH 10 | 0.143 | 0.152 | 2.20 | 0.21 | 0.34±0.66 | Small |
|  | pH 4.5 | 0.0678 | 0.0455 | 3.22 | 0.27 | 0.82±0.68 | Large |
|  | Salt | 0.0658 | 0.0939 | 4.19 | 0.29 | 0.56±0.67 | Medium |
|  | H<sub>2</sub>O<sub>2</sub> | 0.161 | 0.193 | 6.54 | 0.23 | 1.22±0.71 | Large |
| Fitness in<br>complex<br>environments | No. 51 | 0.037 | 0.054 | 4.05 | 0.38 | 0.75±0.68 | Medium |
|  | No. 54 | 0.047 | 0.058 | 1.16 | 0.71 | 0.42±0.66 | Small |
|  | No. 68 | 0.064 | 0.076 | 18.65 | 0.07 | 0.9±0.69 | Large |
|  | No. 49 | 0.126 | 0.155 | 5.67 | 0.32 | 0.79±0.68 | Medium |
|  | No. 22 | 0.078 | 0.084 | 0.15 | 0.71 | 0.29±0.66 | Small |
|  | No. 13 | 0.095 | 0.102 | 0.23 | 0.88 | 0.34±0.66 | Small |
| Fitness in<br>novel<br>environments | Cobalt | 0.019 | 0.079 | 308.37 | 2.4E-04 | 2.71±0.90 | Large |
|  | Norfloxacin | 0.024 | 0.070 | 68.02 | 1.3E-03 | 3.34±1.01 | Large |
|  | EtBr | 0.037 | 0.055 | 111.88 | 2.3E-03 | 1.14±0.70 | Large |
|  | Zinc | 0.030 | 0.029 | 0.63 | 0.47 | 0.20±0.65 | Small |
| Fitness post<br>selection in<br>deteriorating<br>environments | Cobalt | 0.083 | 0.093 | 9.56 | 0.03 | 2.08±1.14 | Large |
|  | Norfloxacin | 0.092 | 0.104 | 17.81 | 0.03 | 1.91±1.11 | Large |
|  | Zinc | 0.054 | 0.060 | 13.15 | 0.04 | 1.20±1.00 | Large |

**Fig 2:**
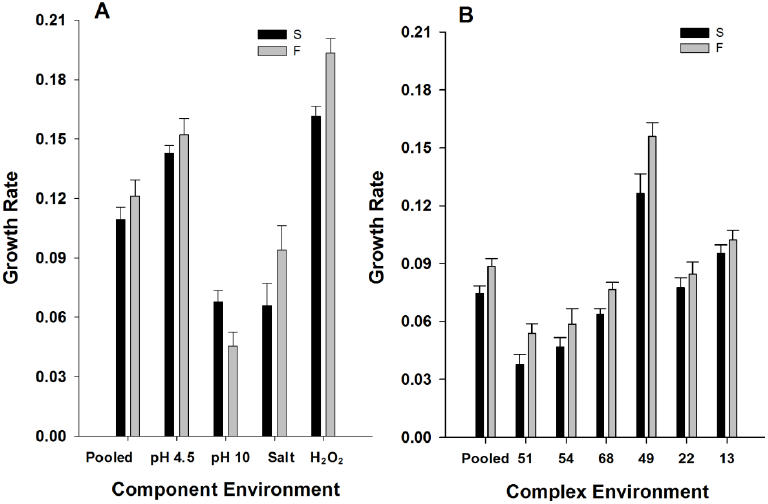
Mean (±SE) fitness in component and complex environments. **A.** Component environments. In the first comparison of the pooled means over all four component environments, the selected (F) populations show significantly higher fitness than the control (S) populations. The next four comparisons are for the individual component environments with no significant difference (individual ANOVAs) in the fitness of F and S populations. **B**. Complex environments. There was no significant difference between the fitness of the F and S populations, either pooled over the six complex environments, or separately in each environment. Fitness was measured as maximum slope of the growth trajectory over 24 hours. * denotes *p* < 0.05.

#### 3.1.2 Fitness under complex environments

The pattern of fitness differences under complex environments was similar to those under component environments. When data from all six complex environments were analyzed together, there was no significant difference between the growth rates of F and S populations (F_1,4_ = 2.37, p = 0.199) and the effect size was small (d = 0.34, Fig 2B, Table 1). When analyzed separately, the average maximum growth rates of the F populations were always larger than those of S populations (Fig 2B), although none of the differences were significant at the 0.05 level, and only one effect size was large (Table 2). Overall, this suggests that the F populations did not show greater fitness in the complex environments.

#### 3.1.3 Fitness under novel environments

When analyzed together, the F populations had significantly higher maximum growth rates under the four novel environments that we investigated (F_1,4_ = 1320.48, *p* = 3.4E-06, Table 1, Fig 3) and the effect size of this difference was large (d = 1.45, Table 1). As with component environments, there was a significant effect of the novel environment (F_3,12_ = 20.38, p = 5.3E-05) and a significant novel environment × selection interaction (F_3,12_ = 48.90, p = 5.3E-07). When analyzed separately, F had significantly greater growth rate than S in three (Cobalt, Norfloxacin, Ethidium bromide) of the four novel environments and the effect size in each of these cases was large (Table 2, Fig 3). In the case of zinc, F still had a higher growth rate than S, although the differences were not statistically significant and the effect sizes were small. However, crucially, the S populations never had a larger growth rate than the F populations. This suggests that the F populations have evolved to counter at least three novel environments without becoming worse than the controls in at least one other novel environment.

**Fig 3:**
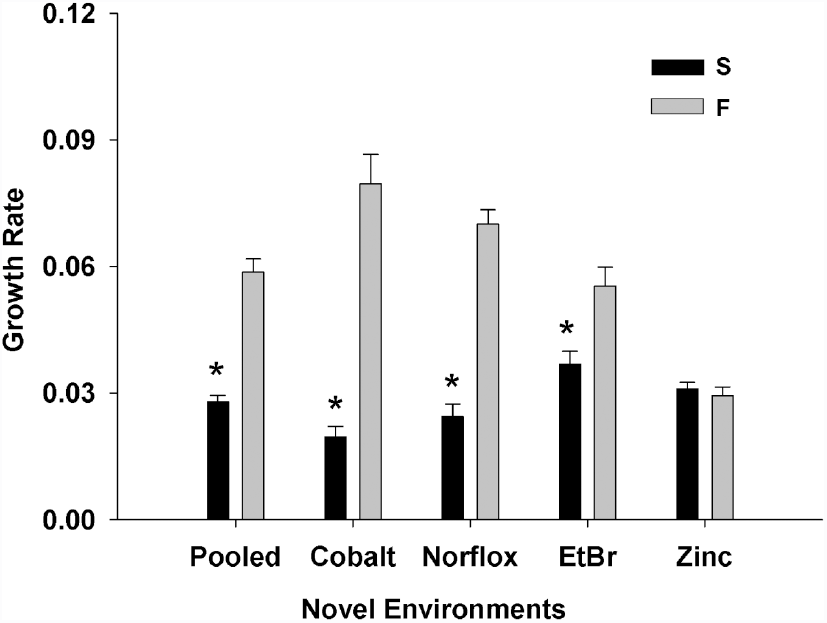
Mean (±SE) fitness in novel environments. Pooled over the four novel environments, F populations show significantly higher fitness than S populations. When compared separately for each novel environment, F populations show significantly higher fitness in cobalt, norfloxacin and ethidium bromide and similar fitness in zinc. * denotes *p* < 0.05 (after Holm-Šidàk correction in the case of comparisons under individual environments).

#### 3.1.4 Fitness after selection under deteriorating novel environments

After directional selection for ∼50 generations in three novel environments, F populations had significantly higher growth rates compared to the S and a medium effect size of the difference (F_1,4_ = 24.68, p = 0.008; *d* = 0.51, Table 1, Fig 4). When analyzed separately, the difference in growth rates in all three novel environments were significant after Holm-Šidàk correction (*p* = 0.036, *p* = 0.039, *p* = 0.043 for Cobalt, Norfloxacin and Zinc respectively, Table 2) and the effect sizes of the differences were large (Table 2).

**Fig 4:**
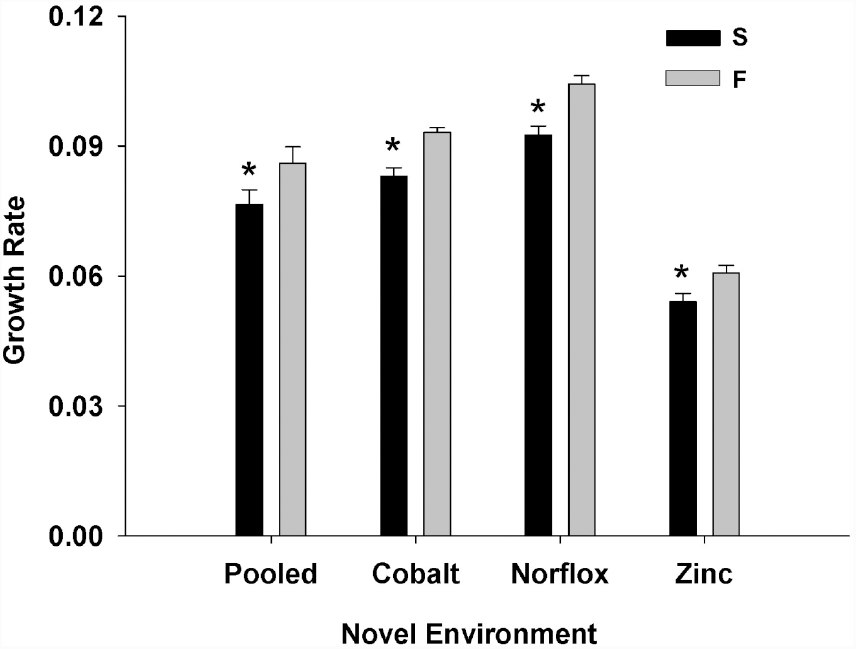
Mean (±SE) fitness in novel environments after facing deteriorating environments. Pooled over the three novel environments, populations selected under fluctuating environments (F) show significantly higher fitness than the control (S) populations. When compared separately for each novel environment, the F populations still had significantly higher growth rate than the S populations. This indicates that the fitness advantages of the F populations were retained after ∼50 generations of directional selection. * denotes *p* < 0.05 (after Holm-Šidàk correction in the case of comparisons under individual environments).

### 3.2 Mechanisms of adaptation

#### 3.2.1 Mutation rates

Although the F populations had higher mutation rates than the S with a large effect size (*d* = 1.65, Table 1), this difference was not statistically significant (F_1,4_ = 4.112, *p* = 0.11; Table 1, Fig 5A). More importantly, the mutation rates were of the order of 10^−8^ to 10^−9^ which suggests that the ability of the F populations to face novel environments is not attributable to the evolution of hypermutators (i.e. alleles that cause 100-1000 fold increase in mutation rate) (Denamur & Matic, 2006).

**Fig 5:**
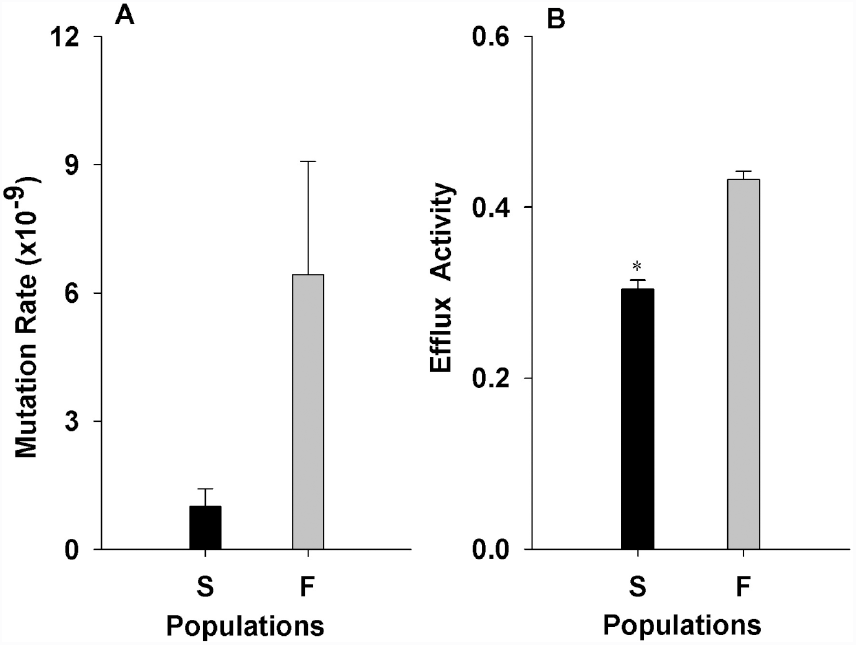
Mechanisms of adaptation. **A.** Mean (±SE) mutation rate for S and F populations estimated in the Rifampicin background. Although the F populations had slightly increased mutation rates than the S populations, the difference was not statistically significant. More importantly, the mutation rates are of the order of 10^−9^, which indicates that hypermutators did not evolve in either of the populations. **B.** Mean (±SE) energy dependent efflux in S and F populations. F populations had significantly greater efflux activity than the S populations, which might have led to their greater fitness in novel environments.

#### 3.2.2 Efflux rates

The F populations had significantly higher activity of the ATP dependent efflux pumps with a large effect size of the difference (*F*_1,4_= 34.65, *p* = 0.004; *d* = 3.79, Table 1, Fig 5B). This indicates the evolution of an increased rate of efflux of toxic materials from inside the cells, which can explain the ability of the F populations to maintain a higher growth rate in novel environments.

The observed increase in efflux activity could have been either due to exposure to unpredictable fluctuations in the environments, or merely due to the component stresses themselves. To distinguish between these two possibilities, we compared the efflux activities of populations directionally selected under the four component stresses (acidic pH, basic pH, salt and H2O2). When analyzed together, there was no main effect of selection in the ANOVA (F_4,10_= 0.577, *p* = 0.686) suggesting that efflux activity did not differ significantly among the populations subjected to the four stresses and those evolved under NB. When we analyzed the efflux activities separately for each stress, none of the differences turned out to be statistically significant at the 0.05 level (see SOM S6). In terms of the effect size of the difference, acidic pH (d = 0.29) and basic pH (d = 0.19) showed small effect sizes. In the case of salt, although the effect size was large (d = 1.13), the salt-selected populations actually had lower efflux activity than the controls. Only in case of H_2_O_2_, the selected populations had a larger efflux activity, and the effect size of the difference was large (d = 1.91). However, given that we failed to get a significant difference either in the pooled ANOVA or the individual one for H_2_O_2_, it is difficult to state that there was a significant change in efflux due to H_2_O_2_. Overall, these results suggest that exposure to unchanging component environments could not have been responsible for the observed increase in efflux activity of the F populations (see section 4.4 for discussion).

## 4 Discussions

### 4.1 No significant adaptation to the component or complex environments

When fluctuations in the selection environment are predictable, organisms often evolve to have higher fitness in the component environments (i.e. conditions experienced during the selection process) (Leroi *et al*., 1994; Turner & Elena, 2000; Hughes *et al*., 2007; Coffey & Vignuzzi, 2011; Alto *et al*., 2013; Puentes-Téllez *et al*., 2013; Condon *et al*., 2014), although some studies report no change in fitness relative to the ancestors or controls (Buckling *et al*., 2003). However, when the selection environment changes unpredictably, fitness in the component environments can increase (Turner & Elena, 2000; Ketola *et al*., 2013b), show no change (Alto et al., 2013), increase w.r.t some life-history traits and decrease w.r.t others (Hallsson and Björklund, 2012) or increase in some of the environments but not all (Hughes et al., 2007). In this study, although we observed an overall statistically significant increase in fitness under component environments (Fig 2), the magnitude of this increase was not biologically meaningful (Table 1), Particularly on short time-scales. In terms of complex environments (i.e. the kind of environments actually faced during selection), the fitness of the selected populations did not differ significantly from the controls and the magnitude of the effect size was low.

There can be multiple (and non-exclusive) reasons for the above observations. Firstly, it is known that the response to selection depends on the rate at which the environment is changing (Venail et al., 2011) and some evolutionary outcomes are possible only when the environments change relatively slowly (Lindsey et al., 2013). In our experiment, the environment changed every 24 hours i.e. every ∼6 generations, which might be too fast for the bacteria to adapt. Secondly, earlier studies on effects of unpredictable environmental fluctuations have mostly been in the context of simple conditions like temperature (Hallsson & Björklund, 2012; Alto et al., 2013; Ketola *et al*., 2013b) or pH (Hughes et al., 2007). However, our study looked at unpredictable combinations of multiple values of three conditions (pH, salt and H_2_O_2_) leading to a total of 72 possible environments, out of which 24 were actually experienced during the course of this study (SOM S1). This means that, on an average, our bacteria were forced to adapt to a novel combination almost every day. When the direction and target of selection changes stochastically, alleles that experience positive selection in a given environment might end up being neutral or negatively selected in the next environment. If the changes in environment happen sufficiently fast (as was the case in our populations), alleles with positive effects on fitness may not get sufficient time to get fixed before the environment changes. As a result of this, at least in the short run, a population might keep on evolving continuously, without really improving in terms of fitness in any of the component environments.

Another reason for the insignificant change in growth rate in component and complex environments might be the relatively short duration of the selection experiment (30 days, ∼170 generations). This appears to contradict a recent study demonstrating that within three weeks of selecting *Serratia* populations under randomly fluctuating temperatures, the population growth rates can increase significantly (Ketola et al., 2013b). However, when we estimated the effect sizes from the fitness data of the previous study (Ketola et al., 2013a), it was found that the Cohen's *d* for the differences in fitness of the control and the selected were small to medium (SOM S7) and never large. It is evidently difficult to compare effect sizes across model organisms. However, if we adopt the same criteria for interpreting Cohen’s *d* statistics for both studies, then the conclusions are identical: selection for short durations leads to statistically significant increase in fitness under component environments, Interestingly, when *E. coli* are subjected to long term selection (∼2000 generations), the magnitude of increase in fitness under unpredictable environments is typically less than that under predictably fluctuating environments (Hughes et al., 2007). This suggests that somehow, stochastic fluctuations hinder the evolutionary increase of fitness more than deterministic fluctuations, which is consistent with similar experiments on viruses (Alto et al., 2013).

To summarize, given the fast rate of change of environment, large number of possible states that selection environment could take and the relatively short duration of the selection experiment, it is not surprising that the F populations in general failed to increase fitness in the component environments. By that line of reasoning, there was no reason to expect any changes in fitness under novel environments either. However, that was not the case.

### 4.2 Short-term and long-term increase in fitness under novel environments

When exposed to novel environments, the F populations had significantly higher fitness in three of the four environments and suffered no disadvantage compared to the controls in the remaining one (Fig 3, Table 2). This suggests that selection under fluctuating environments can increase the ability of populations to face completely novel stresses, which agrees with the results of a previous study (Ketola et al., 2013b). This is also consistent with observations from the literature on invasive species that organisms inhabiting disturbed habitats are often able to cope better with novel environments, thus becoming better invaders (reviewed in Lee & Gelembiuk, 2008).

The simplest explanation for the better performance of the F populations under novel environment could be the presence of cross-tolerance / correlated response to selection for pH, salt and H_2_O_2_. However, we failed to locate any study in the literature that has observed an increased growth rate under any of our novel stresses as a result of selection for any of the three stresses that we used as components for the fluctuating environment. This is perhaps not surprising since the four novel stresses that we used had very different modes of action. For example, while norfloxacin is a DNA gyrase and topoisomerase IV inhibitor (Drlica & Zhao, 1997), ethidium bromide is a DNA intercalating agent which can cause frame-shift mutations (Singer et al., 1999). Cobalt affects the activity of iron-sulfur enzymes (Ranquet et al., 2007) but zinc acts primarily by preventing manganese uptake by cells (McDevitt et al., 2011). Although, in principle, it is possible that the F populations had evolved four different mechanisms to fight each of these stresses through correlated response(s), the likelihood of such an event was low, particularly given the relatively short duration of the selection. While this lack of evidence from the literature cannot be construed as a proof, it at least forced us to explore the possibility of other mechanisms that might be at work to confer a growth advantage to the F populations upon exposure to novel environments.

Some theoretical studies suggest that environmental fluctuations spanning across a few generations (as in our study) can promote the maintenance of genetic variation in a population (Turelli & Barton, 2004), presumably leading to advantages under novel conditions. A different modeling approach predicts that when environments fluctuate rapidly, organisms of intermediate fitness are selected, which can tolerate a multitude of conditions, but none too well (Meyers et al., 2005). Intuitively, it is expected that any mechanism which allows a population to attain higher growth rates upon first exposure to a novel stress, will also be beneficial under directional selection for that stress. This is because a higher growth rate during the initial stages of exposure to a stress allows the bacterial population to undergo more divisions and hence an increased probability of acquiring a suitable beneficial mutation which can then spread in the population. Thus, the F populations were expected to have greater fitness under directional selection in deteriorating environments, which was indeed found to be the case (Fig 4). After ∼50 generations, the F populations retained an overall higher growth rate than the S populations in all three novel environments (Table 2). However, the magnitude of difference in fitness between F and S populations reduced after 50 generations of directional selection (*cf* Fig 3 and Fig 4). This observation is consistent with a scenario where the selected populations have more standing genetic variation but do not have an increased mutation rate. Greater standing genetic variation makes the existence of a favorable mutation in the population more likely, thus providing an early growth advantage. On the other hand, an increased mutation rate in the F populations is expected to increase the difference in fitness vis-à-vis the controls, at least in the short run (Wagner, 1981), which was not found to be the case.

### 4.3 No significant change in mutation rates

Fluctuating environment are expected to favor an increase in the rate of generation of genetic variation through an increase in mutation rates of the populations (Leigh Jr, 1970; Ishii *et al*., 1989). Although the average mutation rate of our F populations was larger than that of the S (Fig 5A), the difference was not statistically significant. There can be multiple (not mutually exclusive) reasons for this observation. Firstly, ∼170 generations of selection might have been insufficient for the mutation rates of the selected populations to have diverged enough to be statistically distinguishable from those of the controls. The fact that the magnitude of the difference is ∼6 folds and the effect size of the differences is large (Table 1) is consistent with the notion that mutation rates in the selected populations are increasing. Secondly, constitutively-expressed mutator alleles are typically thought to spread in a population by hitch-hiking with a beneficial allele (Sniegowski *et al*., 1997; Taddei *et al*., 1997; Gentile *et al*., 2011), although see (Torres-Barceló et al., 2013). However, as in our experiments, if the direction of selection keeps on changing very often, then it is unlikely that a particular mutation would be favorable for long. This will also make it difficult for an attached mutator allele to hitch-hike to fixation with the selected mutation.

Thus, although it is difficult to pin-point the reason, the chief result here is that the difference between the mutation rates of the selected and the control populations was relatively small and not statistically significant. In other words, there is no evidence to suggest that fluctuating environments select for large increases in mutation rate. Coupled with the relatively short period of selection (~170 generations) it is unlikely that the observed growth rate differences in the novel environments are due to the selected populations generating genetic variation faster than the control populations. This prompted us to look at another class of mechanisms that can possibly lead to broad-based stress resistance in bacteria, namely, multidrug efflux pumps.

### 4.4 Evolution of increased efflux activity

Over the last two decades, bacteria have become resistant to a large number of antibiotics due to, *inter alia*, the action of a variety of multidrug efflux pumps (MEPs) (Li & Nikaido, 2009; Nikaido & Pagès, 2012). As the name suggests, MEPs are protein systems that throw out a wide range of antibiotics and biocides from the cells and can also play a role in combating environmental stresses like bile salts (Thanassi et al., 1997) or organic solvents (Fernandes et al., 2003). Our results indicate that the F populations have evolved significantly higher efflux activity as compared to the controls (Fig 5B). Increased activity of MEPs can lead to resistance to norfloxacin (Morita *et al.,* 1998; Nishino *et al*., 2009) and ethidium bromide (Ma *et al*., 1993; Nishino *et al*., 2009) in *E. coli*, both of which were observed here (Fig 3). Multiple drug resistance is known to be associated with higher MEP activity in different kinds of bacteria (Li & Nikaido, 2009; Nishino *et al*., 2009; Nikaido & Pagès, 2012 and references therein) including *E coli* (Pena-Miller et al., 2013). However, to the best of our knowledge, this is the first laboratory experimental evolution study in *E. coli* that reports the evolution of increased energy-dependent MEP activity and the concomitant change in growth rate in the presence of an antibiotic (norfloxacin) and a mutagen/biocide (ethidium bromide), after an exposure to fluctuating complex environments.

An increased efflux activity could have evolved in our F populations in several ways. Firstly, it could have been accidentally fixed in the population due to genetic drift. However, the chances of such an event are low due to the relatively large population sizes involved in bacterial systems and the observation that the increase in efflux happened in all three replicate F populations. Secondly, increased efflux might have also evolved as a result of a direct or a correlated response to selection. Prima facie, this appears counter-intuitive because multi-drug efflux pumps are often studied in the context of drugs or biocides, although of late they have been shown to confer resistance to components of the *E. coli* environment like bile salts (Thanassi et al., 1997) or steroid hormones (Elkins & Mullis, 2006). Since none of these stressors were present during the selection process, it is hard to see how increased efflux activity could have been directly selected for. This inconsistency gets resolved if we take into account the suggestion made by some authors that MEPs have more functions than removing drugs, including clearing out different kinds of secondary metabolites (Poole, 2005) and virulence (Piddock, 2006). Recently it has been shown that a MEP called *norM* can reduce the intracellular levels of reactive oxygen species and increase the ability of *E. coli* cells to survive H2O2 (Guelfo et al., 2010). Since H_2_O_2_ was a constituent of our fluctuating environment, all else being equal, any change leading to increased levels of *norM* should be favored. Incidentally, among all the component environments, the effect size of the difference between S and H was the maximum for H_2_O_2_ (Table 2). This is again consistent with the expectation that resistance to H_2_O_2_ has experienced positive selection. Moreover, the resistance to low pH in *E. coli* is mediated by a regulator called GadX (Ma et al., 2002) which has been shown to elevate the levels of another MEP called mdtEF (Nishino et al., 2008). Finally, the levels of acrAB (a well-studied MEP), are increased by higher concentrations of NaCl (Ma et al., 1995). Thus it was possible that somehow, the efflux activity had evolved due to a direct response to the component environments, without any role of the unpredictable fluctuations.

When exposed to constant stress (acidic pH or basic pH or salt or H_2_O_2_), replicate bacterial populations failed to evolve significantly larger efflux activity, compared to populations evolving in constant environment (NB). Although 40 generations seems a relatively short duration, it should be noted here that the F populations experienced a different environment every ~5.64 generations whereas the populations selected for the constant stresses did not. Thus, if indeed the component environments were selecting for increased efflux activity, the latter set of populations were under much stronger directional selection for the same and hence were expected to show appreciable response to selection. Although the H2O2 selected populations had somewhat greater mean efflux activity compared to the controls (d = 1.91), the lack of statistical significance in the pooled or the individual ANOVA prevents us from concluding that exposure to H_2_O_2_ alone led to appreciable change in efflux activity during this span of time. It is possible that further exposure to H_2_O_2_ might have led to greater changes in efflux activity in the constant stress selected populations. However, given that the F populations never experienced 7 or more continuous runs of exposure to H_2_O_2_, the evolution of efflux activity under such a scenario has no bearing on the results of the present study.

It should be noted here that there is also a possibility of an interaction between unpredictable fluctuations and the component environments in terms of evolution of efflux activity. In other words, some of the component environments (and not others) when fluctuated unpredictably, might have led to the evolution of increased efflux activities. It is also possible that the evolution of increased efflux activity is due to the complex environments (i.e. the combinations of stresses) or an interaction of the complex environments with unpredictable fluctuations. Given that our F populations experienced the complex environments randomly and not every complex environment is likely to select for increased efflux, it is not intuitively obvious that complex environments, by themselves, would lead to the evolution of efflux activity. However, the possibility for the same cannot be completely ruled out, thus forming a fruitful avenue for future research.

## 5 Conclusions

Since bacterial MEPs are often broad-spectrum (Nikaido & Pagès, 2012), it is likely that our F populations would show greater fitness against several other kinds of stresses, including antibiotics, and this advantage might be maintained even after directional selection in the novel environments. This suggests that these populations have evolved greater evolvability, i.e. ability to respond to selection pressure (Colegrave & Collins, 2008). Generally, evolvability is thought to increase through elevated mutational supply (Taddei et al., 1997), or increased rates of genetic exchange (Colegrave & Collins, 2008) or increase in the neutral space of the selected populations (Wagner, 2005). Here we show that evolvability can also evolve in bacteria through the evolution of broad-spectrum stress resistance mechanisms such as MEPs, with potentially serious consequences for the society. Over the last three decades, the day-to-day variability in climate has increased across the globe (Medvigy & Beaulieu, 2012). Our results indicate that unpredictable environmental fluctuations at this time scale can lead to the evolution of enhanced efflux activity, which in turn can foster multi-drug resistance in bacteria. This is a scary scenario from a public health perspective, and no amount of judiciousness in drug-use schedules can prevent this kind of evolution. Moreover, due to the extremely broad substrate ranges for such pumps, it is hard to predict what other traits might evolve when bacteria are subjected to such stochastic fluctuations, particularly under natural conditions. Taking a broader perspective, it would be interesting to know how more complex multi-cellular organisms respond to increased environmental unpredictability. These questions directly relate to the immediate future of the living world, and therefore, merit further investigation.

## Acknowledgements

We thank Amitabh Joshi, Milind Watve, LS Shashidhara, Shashikant Acharya and Neelesh Dahanukar for several illuminating discussions. SK was supported by a Senior Research Fellowship from Council of Scientific and Industrial Research, Govt. of India, while AA thanks Indian Academy of Sciences, for financial assistance through a Summer Research Fellowship Program. This project was supported by a grant from Department of Biotechnology, Government of India and internal funding from Indian Institute of Science Education and Research, Pune. The authors declare no conflict of interests.

## SUPPORTING INFORMATION

### S1 Details of the selection regime faced by F populations

**Table S1.**

S1 – Details of the selection regime faced by F populations
| Transfer # | Combination # | pH | Salt (g%) | H<sub>2</sub>O<sub>2</sub> (M) |
| --- | --- | --- | --- | --- |
| 1 | 46 | 9.5 | 2 | 0 |
| 2 | 54 | 9.5 | 4 | 0 |
| 3 | 31 | 7 | 5 | 0.014 |
| 4 | 64 | 4.5 | 5 | 0 |
| 5 | 31 | 7 | 5 | 0.014 |
| 6 | 30 | 7 | 4.5 | 0.014 |
| 7 | 26 | 7 | 1 | 0.014 |
| 8 | 13 | 7 | 2 | 0.012 |
| 9 | 10 | 9.5 | 0.5 | 0.01 |
| 10 | 68 | 8 | 5 | 0 |
| 11 | 51 | 10 | 3 | 0 |
| 12 | 20 | 6 | 0.5 | 0.012 |
| 13 | 22 | 8 | 0.5 | 0.012 |
| 14 | 15 | 7 | 4 | 0.012 |
| 15 | 55 | 10 | 4 | 0 |
| 16 | 1 | 7 | 5 | 0 |
| 17 | 16 | 7 | 4.5 | 0.012 |
| 18 | 53 | 5 | 4 | 0 |
| 19 | 71 | 10 | 5 | 0 |
| 20 | 54 | 9.5 | 4 | 0 |
| 21 | 8 | 4.5 | 0.5 | 0.01 |
| 22 | 29 | 7 | 4 | 0.014 |
| 23 | 38 | 9.5 | 0.5 | 0.014 |
| 24 | 37 | 9 | 0.5 | 0.014 |
| 25 | 32 | 4.5 | 0.5 | 0.014 |
| 26 | 68 | 8 | 5 | 0 |
| 27 | 15 | 7 | 4 | 0.012 |
| 28 | 49 | 5 | 3 | 0 |
| 29 | 49 | 5 | 3 | 0 |
| 30 | 51 | 10 | 3 | 0 |

Composition of Nutrient broth –

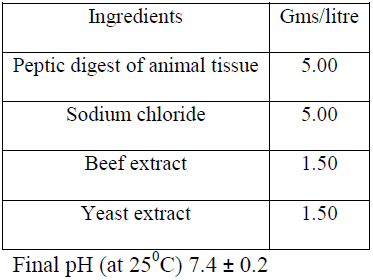

| Ingredients | Gms/litre |
| --- | --- |
| Peptic digest of animal tissue | 5.00 |
| Sodium chloride | 5.00 |
| Beef extract | 1.50 |
| Yeast extract | 1.50 |

Final pH (at 25<sup>0</sup>C) 7.4 ± 0.2

Component space for fluctuating selection regime –

Salt (g%) - 1, 2, 3, 4, 4.5, 5

pH - 4.5, 5, 6, 7, 8, 9, 9.5, 10

H_2_O_2_ (M) – 0, 0.01, 0.012, 0.014

### S2 Novel, component and complex environments used for fitness assays after fluctuating selection

**Table S2.**

**S2 – Novel, component and complex environments used for fitness assays after fluctuating selection**
|  | **Assay Environment** | **Concentration** |
| --- | --- | --- |
| Novel environments | Cobalt chloride | 14.2 mg% |
|  | Zinc sulfate | 18.4 mg% |
|  | Ethidium bromide | 2.5 mg% |
|  | Norfloxacin | 0.004 mg% |
| Component environments | Salt | 5 g% |
|  | Acidic pH | 4.5 |
|  | Basic pH | 10 |
|  | Hydrogen peroxide | 0.058 M |
| Complex environments | #51 | pH 10 + salt 3g% + 0 H<sub>2</sub>O<sub>2</sub> |
|  | #54 | pH 9.5 + salt 4g% + 0 H<sub>2</sub>O<sub>2</sub> |
|  | #68 | pH 8 + salt 5g% + 0 H<sub>2</sub>O<sub>2</sub> |
|  | #49 | pH 5 + salt 3g% + 0 H<sub>2</sub>O<sub>2</sub> |
|  | #22 | pH 8 + salt 0.5g% + 0.01M H<sub>2</sub>O<sub>2</sub> |
|  | #13 | pH 7 + salt 2g% + 0.01M H<sub>2</sub>O<sub>2</sub> |

### S3 Selection under deteriorating environment

**Table S3.**

S3 - Selection under deteriorating environment
| Environment | Concentrations |  |  |
| --- | --- | --- | --- |
|  | Day I | DayVIII | Step size |
| Cobalt | 6 mg% | 13 mg% | 1 mg% |
| Zinc | 5 mg% | 12 mg% | 1 mg% |
| Norfloxacin | 0.4 µg% | 1.1 µg% | 0.1 µg% |

### S4 Protocol for estimation of mutation rate

We used the method suggested by Crane et al (Crane et al. 1996) for estimating mutation rates. One control population and one selected population were processed with a given batch of media. Glycerol stocks of control S and F populations were revived in NB for 18 hours. The revived culture was then diluted 10 fold. Eleven replicate flasks, each containing 50 ml of nutrient broth, were inoculated with 10 μl each of this diluted suspension. These replicate cultures were then allowed to grow for another 18 hours and then plated on nutrient agar with and without 100 micrograms/ml Rifampicin (Torres-Barceló et al. 2013) for estimating mutant counts and total numbers respectively. Colonies were counted manually, after 24 hours of incubation at 37ºC. Median estimator (λ_med_) was estimated using following formula (Jones 1994):

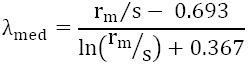

where r_m_ = Number of mutants found in the median culture
s = Proportion of culture plated

Mutation rate, μ = λ_med_/N
where, N = Total number of living bacteria in 50 ml culture (as counted on Nutrient Agar).

### S5 Protocol for estimation of energy dependent efflux activity

For characterization of efflux properties of our populations, we modified an existing protocol (Webber and Coldham 2010) for fluorescent-based estimation of active efflux in Gram negative bacteria. Glycerol stocks of S and F populations were revived for 18 to 20 hours in NB. OD_600_ was adjusted, by diluting with NB, in the range of 0.03 to 0.06 OD units on Nanodrop (Thermo scientific 2000c). 2 ml of the OD_600_ adjusted cultures were then centrifuged, supernatant was discarded and the pellet was re-suspended in the PBS buffer (pH 7.4). Part of each suspension was then boiled at 60°C for 10 minutes to be used as positive controls. The live cells and corresponding positive controls were then loaded in 96 welled plates in triplicates (168 μl and 180 μl respectively). 20 μl of Bis-benzimide (Excitation λ 355 nm and Emission λ 465 nm) was then added to all the wells. Bis benzimide is a small molecule which can easily enter the cell and fluoresce after intercalating with DNA. Live cells were also supplied with glucose since we wanted to look at the ATP dependent active efflux (8 μl of 1% glucose solution). The total volume of each well with live cell was 196 μl and with dead cells was 200 μl.

Fluorescence was measured on a plate reader (Tecan Infinite M200 Pro) at 37°C for forty minutes. By 35 minutes the fluorescence levels in all wells reached their steady state. The level of fluorescence at 35 minutes was taken as the total fluorescence. The reader was then paused and 4 μl of Carbonyl Cyanide m-Chlorophenylhydrazone (C2759 Sigma) was added to all the wells with live cells. Carbonyl Cyanide m-Chlorophenylhydrazone (CCCP) is a non-specific inhibitor of active efflux in Gram negative bacteria (Webber and Coldham 2010). Fluorescence measurement was continued for another 30 minutes for steady state to reach and the reading at 70^th^ minute was taken as fluorescence without efflux.

The contribution of efflux was measured as –

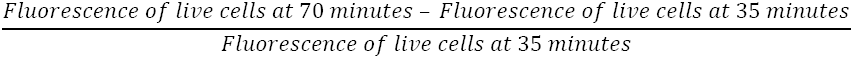

### S6 Efflux activity under selection for adaptation to constant environment

Median values for each of the stress parameter (Salt 4g%, pH5, pH9.5 and 0.012M H_2_O_2_) were chosen for the selection in constant environment. *Escherichia coli* (strain NCIM 5547) was revived overnight in Nutrient Broth. This revived culture was used to initiate three replicate populations in each of the selection environments and Nutrient Broth, a total of 15 populations. Culturing conditions and transfer volume was as mentioned in section *2.1.1*. The selection lasted for seven days (i.e ~40 generations), without any extinction events, after which the populations were stored as glycerol stocks. These stocks were then used for efflux measurement as per section SOM-S5.

#### Statistical Analysis

The average of the three efflux measurements for each population was used for analyzing all the environments together. The pooled data was analyzed using 1-way ANOVA where Selection (5 levels: Salt, pH5, pH9.5, H_2_O_2_ and NB) was a fixed factor. For analyzing each stress separately, we performed four separate 2-way mixed model ANOVAs where selection (2 levels: selected and control) and replicates (3 levels, nested in selection) were treated as fixed and random factors respectively.

#### Results

Efflux did not differ across environments (*F*_4,10_= 0.577, *p* = 0.686). Results of individual ANOVA, summarized in the following table, show no difference in the energy dependent efflux of control and any of the selected populations when analyzed separately.

**Table S6.**

| Selection environment | Mean Control | Mean Selected | ANOVA F(1,4) | ANOVA *p* values | Effect Size±95% CI | Inference |
| --- | --- | --- | --- | --- | --- | --- |
| Salt | 0.648 | 0.49 | 1.52 | 0.285 | 1.13±0.99 | Large |
| pH 5 | 0.551 | 0.507 | 0.1 | 0.765 | 0.29±0.93 | Small |
| pH 9.5 | 0.59 | 0.622 | 0.04 | 0.851 | 0.19±0.93 | Small |
| H<sub>2</sub>O<sub>2</sub> | 0.458 | 0.626 | 4.74 | 0.095 | 1.91±1.11 | Large |

Note that no Holm-Šidák correction was done since even the lowest *p*-value was not significant at the 0.05 level.

### S7 Effect size computed for the growth rates and yield measured in the three temperature environments from Figure 1 of (Ketola et al. 2013). Means and ANOVA p-values are as reported in that paper

**Table S7.**

|  | Assay Environment | Mean C | Mean F | ANOVA *p* values | Effect Size±95% CI | Inference |
| --- | --- | --- | --- | --- | --- | --- |
| Growth rate | 24<sup>0</sup>C | 0.31 | 0.34 | <0.001 | 0.43±0.25 | Small |
|  | 31<sup>0</sup>C | 0.46 | 0.51 | <0.001 | 0.53±0.25 | Medium |
|  | 38<sup>0</sup>C | 0.37 | 0.41 | <0.001 | 0.61±0.25 | Medium |
| Yield | 24<sup>0</sup>C | 0.086 | 0.089 | <0.001 | 0.21±0.25 | Small |
|  | 31<sup>0</sup>C | 0.1 | 0.11 | 0.478 | 0.12±0.25 | Small |
|  | 38<sup>0</sup>C | 0.075 | 0.079 | <0.001 | 0.45±0.25 | Small |

